# A polygenic architecture with conditionally neutral effects underlies ecological differentiation in *Silene*

**DOI:** 10.1101/2021.07.06.451304

**Authors:** Susanne Gramlich, Xiaodong Liu, Adrien Favre, C. Alex Buerkle, Sophie Karrenberg

**Affiliations:** Department of Ecology and Genetics, Uppsala University, Norbyvägen 18D, 75267 Uppsala, Sweden; The Bioinformatics Center, Department of Biology, University of Copenhagen, Ole Maaløes Vej 5, 2200, Copenhagen, Denmark; Senckenberg Research Institute and Natural History Museum, Senckenberganlage 25, 60325 Frankfurt/Main, Germany; Department of Botany, University of Wyoming, 1000 E. University Ave. Laramie, WY82071, USA

**Keywords:** adaptation, Bayesian Sparse Linear Mixed Models (BSLMM), conditional neutrality, ddRAD-Seq, reproductive isolation, speciation, *Silene*

## Abstract

- Ecological differentiation can drive speciation but it is unclear how the genetic architecture of habitat-dependent fitness contributes to lineage divergence. We investigated the genetic architecture of cumulative flowering, a fitness component, in second-generation hybrids between *Silene dioica* and *S. latifolia* transplanted into the natural habitat of each species.
- We used reduced-representation sequencing and Bayesian Sparse Linear Mixed Models (BSLMMs) to analyze the genetic control of cumulative flowering in each habitat.
- Our results point to a polygenic architecture of cumulative flowering. Allelic effects were mostly beneficial or deleterious in one habitat and neutral in the other. The direction of allelic effects was associated with allele frequency differences between the species: positive-effect alleles were often derived from the native species, whereas negative-effect alleles, at other loci, tended to originate from the non-native species.
- We conclude that ecological differentiation is governed and maintained by many loci with small, conditionally neutral effects. Conditional neutrality may result from differences in selection targets in the two habitats and provides hidden variation upon which selection can act. Polygenic architectures of adaptive differentiation are expected to be transient during lineage divergence and may therefore be unrelated to high genetic differentiation at the underlying loci.

## Introduction

Adaptation to different habitats can promote divergence and speciation and can mediate population persistence under changing conditions (Nosil, 2012; Savolainen *et al*., 2013). Evolutionary trajectories toward adaptive differentiation depend on the number and effect sizes of loci controlling fitness, on allelic effects of such loci in alternative habitats and on gene flow (Savolainen *et al*., 2013; Tigano & Friesen, 2016; Kokko *et al*., 2017). However, despite recent progress in the theoretical understanding of adaptation (Yeaman, 2015; Mee & Yeaman, 2019; Booker *et al*., 2021), empirical studies on the genetic control of fitness in natural habitats is still scarce (Wadgymar *et al*., 2017). Here we provide such a study using two ecologically differentiated but hybridizing campions (*Silene*).

Adaptive differentiation is often depicted as a classic cartoon with populations outperforming other populations in their home habitat but not in the foreign habitat (Kawecki & Ebert, 2004). A similar pattern has been proposed to operate at the genetic level: alleles with positive effects on fitness in one habitat have negative effects in the alternative habitat, termed antagonistic pleiotropy (Anderson *et al*., 2011; Savolainen *et al*., 2013). Empirical data, however, suggests that such antagonistic pleiotropy is rare in natural populations, although this could partially be caused by limited statistical power (Anderson *et al*., 2011; Anderson *et al*., 2014; Wadgymar *et al*., 2017). An alternative pattern, conditional neutrality, where alleles have neutral effects in one environment and are under positive or negative selection in the other, may be much more prevalent (Anderson *et al*., 2011; Savolainen *et al*., 2013; Wadgymar *et al*., 2017). In fact, even at the phenotypic level, populations often perform differently in one habitat but not in the other (Leimu & Fischer, 2008). Importantly, loci with antagonistic pleiotropy for fitness in alternative habitats are expected to be maintained by natural selection and contribute to population differentiation even with high gene flow, whereas conditionally neutral loci can only lead to transient allele frequency differences in high-gene flow scenarios (Mitchell-Olds *et al*., 2007; Savolainen *et al*., 2013; Tiffin & Ross-Ibarra, 2014; Mee & Yeaman, 2019; Booker *et al*., 2021). If adaptive allele frequency changes are predominantly transient, even in the presence of consistent selection, prospects will be poor for finding loci underlying adaptive differentiation with commonly used methods for quantifying and comparing genetic differentiation along the genome (i.e., genome scans of differentiation).

A further major determinant of adaptive differentiation concerns the number and effect sizes of the underlying loci. In general, the distribution of effect sizes for complex traits (including fitness) is expected to be exponential (Orr, 1998, 2005) such that adaptation is mainly due to many loci with small effects, while large-effect loci, although not generally unimportant, are rare (Orr, 1998; Orr, 2005; Rockman, 2012; Savolainen *et al*., 2013; Boyle *et al*., 2017; Selby & Willis, 2018). A large number of segregating loci underlying adaptation further promotes transient genetic architectures of adaptation (Yeaman & Whitlock, 2011; Yeaman, 2015; Booker *et al*., 2021). Polygenic or even omnigenic genetic architectures, involving most of the genome, render identification of all individual loci both practically impossible and undesirable (Rockman, 2012). As effect sizes decrease and the number of loci increases, it becomes progressively more difficult to detect individual genotype-phenotype associations and to distinguish them from spurious associations due to linkage. For this reason, promising methods, such as Bayesian Sparse Linear Mixed Models (BSLMMs), identify genotype-phenotype associations of all loci simultaneously rather than individually, assume polygenic genetic architectures and remove effects that can be attributed to linkage disequilibrium (Zhou *et al*., 2013; Gompert *et al*., 2017). The approach has been successfully applied in studies on adaptive divergence, for example in pine (Lind *et al*., 2017) and in *Arabidopsis* (Exposito-Alonso *et al*., 2019).

In this study, we investigate the genetic basis of differential adaptation in two dioecious sister species of *Silene* (Caryophyllaceae), *Silene dioica* (L.) Clairv. and *S. latifolia* Poiret. Demographic models indicate that the two species diverged with gene flow within the last 120 000 years reaching a neutral sequence divergence (Da, autosomes) of 0.0027 and genetic differentiation, F_ST_, of 0.28 (Hu & Filatov, 2015; Liu *et al*., 2020). The pink-flowered *S*. *dioica* occurs in moister and colder habitats like meadows, pastures or forests and occupies a wide range of elevations up to more than 2300 m a. s. l., while the white-flowered *S. latifolia* is found on drier, warmer and more disturbed sites like dry meadows, arable fields or road sites at elevations up to 1000 m a. s. l. (Friedrich, 1979; Karrenberg & Favre, 2008). Ecological differentiation and assortative pollination constitute strong barriers to gene flow in this species pair, but the two species are still fully cross-fertile (Goulson & Jerrim, 1997; Karrenberg *et al*., 2019). Studies in an experimental garden point to a complex, genome wide genetic architecture of traits associated with reproductive isolation, such as flower color, flowering phenology, first-year flowering and specific leaf area in this system (Liu & Karrenberg, 2018).

Here we investigated the genetic architecture of adaptation using recombinant second-generation hybrids (F_2_) from a multi-site field transplant experiment where the two species exhibited strong evidence of habitat adaptation: each species outperformed the other in its own habitat in terms of flowering and survival over four years (Fig. 1, Favre *et al*., 2017). We focused on the following questions: (1) What is the genetic architecture underlying differential habitat adaptation? (2) Is antagonistic pleiotropy or conditional neutrality the predominant pattern when comparing allelic effects across habitats? and (3) Are fitness effects associated with allele frequency differences between the two species; for example, are beneficial alleles more likely to be derived from the native species?

**Figure 1.**
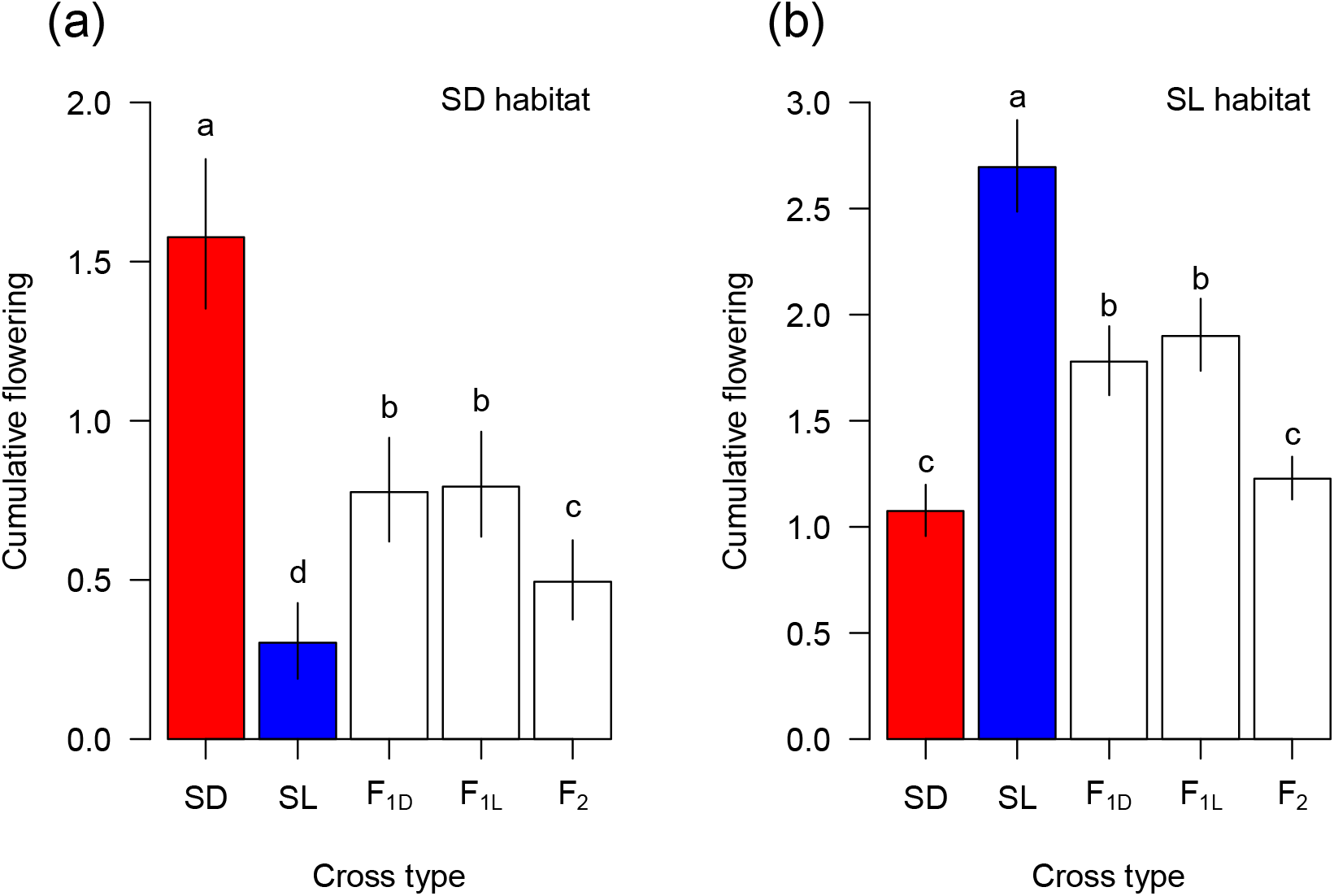
Habitat adaptation in campions (*Silene)*: cumulative flowering (number of times flowered over four years, plus 1 if the plant survived to the end of the experiment), a fitness component, in crosses within each of two campion species, *Silene dioica* (SD) and *S. latifolia* (SL), as well as in their first-generation hybrids (F_1D_ with SD mother, F_1L_, SL mother) and second-generation hybrids (F2) in the habitat of each species, (a) SD habitat and (b) SL habitat; least square means and 95% confidence intervals are given and different letters indicate significant differences between cross types within each habitat. This data is published in Favre et al. (2017 and re-analyzed here for illustration (see text).

## Materials and Methods

### Transplant experiment, measurements and sampling

Second-generation (F_2_) hybrids between *Silene dioica* (L.) Clairv. and *Silene latifolia* Poir., derived from 36 F_0_ individuals of three populations of each species, were transplanted as juveniles into four natural sites, two in each species’s habitat, as part of a larger experiment with six sites (Favre *et al*., 2017, Supporting Information Tables S1, S2, and S3). Sites of *S. dioica* were situated at higher altitudes with a colder climate and shorter growing season as compared to the *S. latifolia* sites (Supporting Information Table S1). Leaf samples were collected prior to transplantation and silica-dried and we selected four of the six sites for this study based on availability and quality of leaf samples. Flowering and survival were assessed over four years and cumulative flowering was calculated as the number of times an individual flowered plus 1 if it survived to the end of the experiment, for detailed analyses of survival and flowering see Favre et al. (2017). We sampled 4-6 F_2_ individuals from each of 18 F_2_ families at each of the four selected sites (298 F_2_ individuals in total), striving to include both high and low fitness individuals from each family and site. To assess allele frequency in the two species we further included samples of 32 F_0_ individuals and used existing data for the remaining four F_0_ individuals (Liu & Karrenberg, 2018, Supporting Information Table S3).

For illustration purposes (Fig. 1), we re-analyzed cumulative flowering in the two species and their first- and second-generation hybrids for the four sites used here, (18-36 families per cross type, 5-20 individuals per family, 5 blocks per site, Favre *et al*. 2017, Supporting Information Table S2). We used linear mixed models of cumulative flowering in each habitat with cross type as a fixed factor and family and block nested in site as random factors using the *lme4* package (Bates *et al*., 2015) for R version 4.02 (R Core Team, 2020). We extracted least square means of cumulative flowering and performed multiple comparisons between cross types within sites with Holm correction of P-values in the *emmeans* package (Lenth, 2020). Cumulative flowering was log(Y+1)-transformed to yield normally distributed residuals and improve model fit. Means and standard errors are reported back-transformed to the original scale.

### DNA extraction and sequencing

We extracted genomic DNA from silica-dried leaf tissue with Qiagen’s DNeasy Plant Mini Kit (Qiagen, Germany) and quantified DNA using a Qubit dsDNA HS fluorometer (Life Technologies, Sweden). Double-digest RAD sequencing (ddRAD-seq) libraries were prepared with EcoRI and TaqαI restriction enzymes as described in Liu and Karrenberg (2018). After enzymatic digestion, DNA fragments were ligated with barcoded adaptors and size-selected to approximately 550 bp (Peterson *et al*., 2012). In total, eight 48-plex libraries were sequenced on an Illumina HiSeq 2500 system at the SNP&SEQ technology platform of SciLifeLab, Uppsala, Sweden using 125-bp paired-end chemistry and two libraries per lane. F_0_ individuals were included in 2 libraries to achieve higher coverage.

### Bioinformatic analysis - processing of raw reads and variant filtering

The total sequencing output was 1,382,838,294 reads for 298 F_2_ individuals (mean with one standard error: 4,656,021 ± 233,639 reads) and 461,127,142 reads for 32 F_0_ individuals (mean: 14,410,223 ±1,093,409 reads); data on the remaining four F_0_ individuals (Supplementary Information Table S3) was available from (Liu & Karrenberg, 2018). We processed the ddRAD-sequence reads following the dDocent pipeline (Puritz et al., 2014). After de-multiplexing of raw reads using stacks 2.0b (Catchen *et al*., 2013) and trimming with *fastp* (Chen *et al*., 2018), we used BWA MEM 0.7.17 with default parameters (Li, 2013) to map reads to ddRAD-seq-generated reference contigs, which were previously assembled from eight deeply sequenced individuals of both species and hybrids and corresponded to 95,040,562 bp in total, corresponding to approximately 3.4% of the *S. latifolia* genome (Liu & Karrenberg, 2018; Liu *et al*., 2020). Only a partial genome sequence (one third of the 2.8 Gbp genome) with short scaffolds (N_50_ = 10,785 bp) is currently available for *S. latifolia* (Krasovec *et al*., 2018).

Variants were called with FreeBayes 1.1.0 (Garrison & Marth, 2012) without population priors using the following parameters: minimum mapping quality 30, minimum base quality 20, maximum complex gap 3, minimum repeat entropy 1, binominal-obs-priors 1, and use-best-n-alleles 10. Variants were filtered following O’Leary *et al*. (2018): First, VCFtools 0.1.15 (Danecek *et al*., 2011) was used to retain SNP sites with a minimum depth of 3, quality of 30, mean depth of 10 and allele count of 3. Secondly, we used *Vcffilter* implemented in vcflib/2017-04-04 (https://github.com/vcflib/vcflib) to retain sites with an allele balance either between 0.25 and 0.75 or lower than 0.01, a quality/depth ratio of > 0.25, and a mapping quality ratio between 0.9 and 1.05. We further used *Vcffilter* to remove loci with differences in read pairing between the alleles or with or excessive read depths (parameters as suggested in O’Leary *et al*., 2018). We reduced the dataset to bi-allelic sites and removed 6 F_2_ individuals with more than 99% missing data. SNPs in perfect linkage disequilibrium (r^2^ = 1) within the F_2_ individuals were removed using PLINK 1.9 (Purcell *et al*., 2007, http://pngu.mgh.harvard.edu/purcell/plink/). The filtered dataset contained 290 F_2_ individuals (*S. dioica*-habitat: 134, *S. latifolia-habitat*: 156) and 89’524 loci. Of these, 42’090 loci with both alleles in both habitats and genotypes for more than 95 F_2_ individuals in each habitat were used for further analyses. These loci had an average read depth of 15.02 ± 0.03 (median 13.83) for the F_2_ individuals, 36.85 ± 0.08 (median: 33.09) for the 32 F_0_ individuals sequenced in this study and 41.39 ± 0.17 (median: 36.25) for the 4 F_0_ individuals from Liu and Karrenberg (2018). We used genotype probabilities ranging from 0 to 2, calculated from genotype likelihoods (https://github.com/visoca/popgenomworkshop-gwas_gemma/tree/master/scripts/bcf2bbgeno.pl, accessed 28 January 2019), rather than called genotypes (Nielsen *et al*., 2011). Genotype probabilities of 0 and 2 denote homozygosity for the reference and alternative allele, respectively, while 1 indicates heterozygosity.

### Genetic structure

We checked for genetic structure in the F_2_ individuals using Principal Components Analysis (PCA) with the *ade4* package (Dray and Dufour, 2007), based on genotype probabilities. For this analysis we reduced the dataset to 220 SNP loci with data available in all 290 F_2_ individuals used in the genetic association analysis. Results did not differ from analyses on more or all loci with missing values replaced by average genotype probabilities.

### Genetic association analysis

We used Bayesian Sparse Linear Mixed Models (BSLMM) in GEMMA 0.98.1 (Zhou *et al*., 2013) to investigate the genetic architecture of cumulative flowering. BSLMMs are a combination of linear mixed models, which assume that every variant has an effect, and Bayesian Variable Selection Regression, which assumes that only a small proportion of the variants has an effect (Zhou *et al*., 2013). BSLMM analyses were performed separately for each habitat using cumulative flowering values standardized within sites and genotype probabilities. A centered relatedness matrix was used as a covariate to take account of the family structure and the wide cross, according to the standard BSLMM method implemented in GEMMA (Zhou *et al*., 2013).

We characterized genetic architectures using estimates of the following hyperparameters: the proportion of phenotypic variance explained by all SNPs in the model (*PVE*), the proportion of *PVE* explained by SNPs with nonzero effects (*PGE*), and the number of SNPs with a measurable effect on the phenotype (n-*γ*). In studies with incomplete genome coverage such as this one, *PVE* can be interpreted as broad sense heritability *H^2^*, whereas the product of *PVE* and *PGE* can be interpreted as narrow-sense heritability *h^2^* (Zhou *et al*., 2013; Gompert *et al*., 2017; Bresadola *et al*., 2019). For each SNP locus, we estimated the posterior inclusion probability (*PIP*, the proportion of iterations in which a SNP had a nonzero effect on phenotypic variation) and the effect size of the alternative allele on cumulative flowering. We report both raw effect estimates 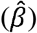, the effect of a locus on the phenotype when it is included in the model, and model-averaged effect estimates 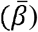 that take into account the posterior inclusion probability of a locus in the models (Zhou *et al*., 2013; Gompert *et al*., 2017). BSLMMs were run five times with 10,000,000 burn-in steps and 40,000,000 iterations with a thinning interval of 10. Convergence of the five runs was checked graphically.

Hyperparameters were estimated after combining posterior distributions across runs; *PIP*, 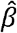 and 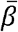 (per locus) were averaged across runs. A threshold of *PIP* > 0.01 was used to identify SNPs with sparse effects (Gompert *et al*., 2013; Comeault *et al*., 2014).

### Allelic effects and allele frequencies

We graphically evaluated whether alleles have universal effects or display antagonistic pleiotropy or conditional neutrality by plotting per-locus effect sizes (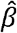 and 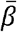) in the *S. dioica* habitat against effect sizes in the *S. latifolia* habitat. Note that *β*-values reflect effect estimates of the alternative allele replacing the reference allele. The *de novo* ddRADseq reference sequences used here contain alleles derived from both species (Liu & Karrenberg, 2018; Liu *et al*., 2020) and do not inform on allelic origin.

To analyze whether allelic origin is associated with fitness effects, we calculated the frequency of the alternative allele (AF_alt_) in the 18 F_0_ individuals of each species (Tables S2 and S3) for loci that had genotypes for > 12 F_0_ individuals per species at individual read depths of at least 6 (36,161 loci). For each SNP locus, we expressed AF_alt_ as the average genotype probability per species divided by 2. We further estimated the allele frequency difference between species (AFD_alt_) as AF_alt_ (*S. dioica*) - AF_alt_ (*S. latifolia*). AFD_alt_ values thus range from −1 (alternative allele fixed in *S. latifolia* F_0_ individuals, and reference allele fixed in *S. dioica* F_0_ individuals) to 1 (alternative allele fixed in *S. dioica* F_0_ individuals, and reference allele fixed in *S. latifolia* F_0_ individuals). We tested whether AFD_alt_ differs between loci with positive and negative effects on cumulative flowering in each habitat using a general permutation test in the R package *coin* (Hothorn et al., 2008).

### Method validation

We assessed the power of BSLMMs to detect genotype - phenotype associations in our data using a simulation approach similar to that in Gompert *et al*. (2017). Phenotypes were simulated on the basis of the observed genetic data in 290 individuals for four combinations of heritability (*h^2^* = 0.2 or *h^2^* = 0.05) and number of functional variants (*N* = 10 or *N* = 50). We simulated a normally distributed trait for which nine individuals of each family were randomly drawn. We reduced the simulated dataset to 150 individuals at random to match the sample size in our data. For each combination of *h^2^* and n 30 sets of simulated phenotypes were used as input for BSLMMs with 10,000,000 burn-in steps and 40,000,000 iterations. For each simulation, *PVE*, *PGE* and *n-γ* were estimated as well as the correlation between the simulated effect size and effect size estimates (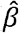 and 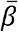).

## Results

### Cumulative flowering and genetic structure

Each species outperformed the other in its own habitat and performed best in its habitat in terms of cumulative flowering, providing strong evidence for habitat adaptation (Fig. 1, Favre *et al*., 2017). F_1_ hybrids were intermediate for cumulative flowering and F_2_ hybrids flowered less often than F_1_ hybrids on average (Fig. 1, Favre *et al*., 2017).

The F_2_ families exhibited extensive genetic variation in the PCA, with clustering of individuals within families but without any other emerging patterns (Supplementary Information Figs. S1 and S2). For most loci, the alternative allele occurred at low frequency in one or both species, but all combinations of allele frequencies between species were found in the data, including loci with high differentiation where the alternative allele was at high frequency in one species and at the same time near absent from the other species (Fig. 2).

**Figure 2.**
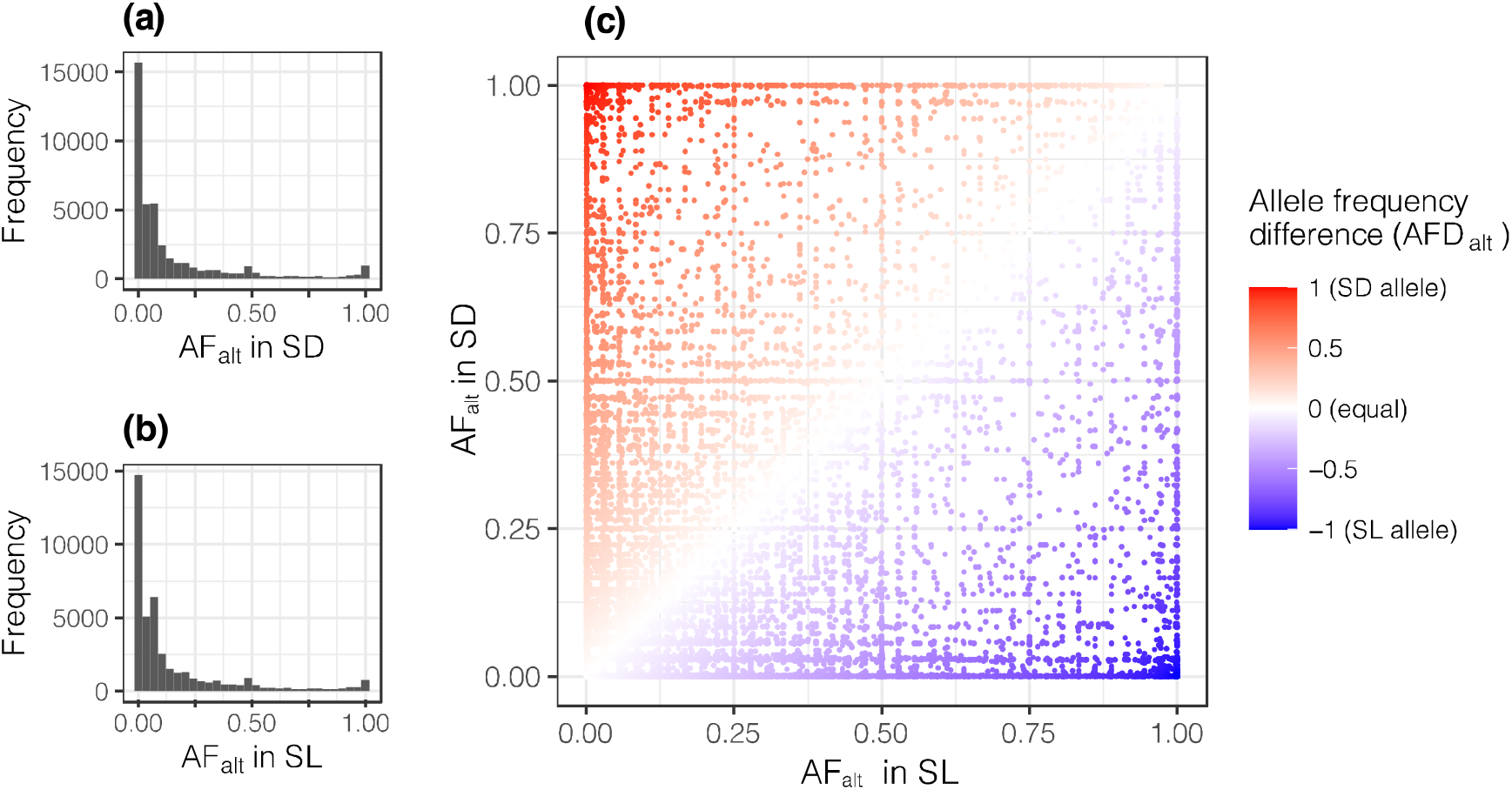
Frequencies of the alternative allele (AF_alt_) in two campion species, *Silene dioica* (SD) and *S. latifolia* (SL) for 42’090 loci used in an association study: **(a)** distribution of alleles frequencies in SD, **(b)** distribution of allele frequencies in SL and **(c)** SD allele frequency (y-axis) against SL allele frequency (x-axis) with allele frequency differences (AFD_alt_) between the two species (SD - SL) indicated as a color gradient from red (allele fixed in SD and absent in SL) to white (equal frequency in both species) to blue (allele fixed in SL and absent in SD).

### Association analysis

The proportion of phenotypic variation explained (*PVE*) by BSLMMs was 0.12 and 0.10 for the *S. dioica* and the *S. latifolia* habitat, respectively (medians of posterior distributions, Table 1, posterior distributions are shown in the Supporting Information Figs. S3 and S4). Less than half of the *PVE* could be attributed to sparse effects, leading to estimates of narrow-sense heritability (*h^2^*; *PGE* × *PVE*) of 0.05 and 0.03 for the *S. dioica* and *S. latifolia* habitat (Table 1). The number of loci with non-zero effects (*n-γ*) was estimated to be 11 in the *S. dioica* habitat and 16 in the *S. latifolia* habitat (Table 1). Median *PIP* for individual SNPs was 0.001 in both habitats and we detected 3 loci with *PIP* > 0.01, one in the *S. dioica* habitat and two in the *S. latifolia* habitat (Supporting information Table S4). *PIP* for individual SNPs could not be summed over genomic windows because no continuous reference genome is available (Krasovec *et al*., 2018) and we used a ddRADseq-generated *de novo* reference in this study (Liu & Karrenberg, 2018; Liu *et al*., 2020).

**Table 1.** Genetic architecture of cumulative flowering, a fitness component, in recombinant F_2_ hybrids between the campions *Silene dioica* and *S. latifolia* in a field transplant experiment at sites of each species; given are medians, means and 90 % equal tail credible intervals (90% cred. int.) of hyperparameters estimated in Bayesian Sparse Linear Mixed Models (BSLMMs): *PVE* (proportion variation explained, broad-sense heritability, *H^2^*), *PGE* (proportion variation explained by measurable effects), *h^2^* (narrow-sense heritability, *PVE* * *PGE*) and *n-γ* (number of loci with measurable effects). Posterior distributions of hyperparameters are shown in Supplementary Figures S3 and S4.

|  | Hyperparameter |  |  |  |
| --- | --- | --- | --- | --- |
| | *PVE* | *PGE* | $h^2$ | $n-\gamma$ |
| *Silene dioica* habitat |  |  |  |  |
| median | 0.117 | 0.411 | 0.037 | 11 |
| mean | 0.136 | 0.434 | 0.060 | 42.2 |
| 90% cred. int. | 0.017 - 0.321 | 0 - 0.945 | 0 - 0.198 | 0 - 205 |
| *Silene latifolia* habitat |  |  |  |  |
| median | 0.096 | 0.411 | 0.029 | 16 |
| mean | 0.124 | 0.431 | 0.054 | 51.4 |
| 90% cred. int. | 0.008 - 0.337 | 0 - 0.938 | 0 - 0.193 | 0 - 225 |

### Allele specific effects in alternative habitats and origin of alleles

Raw effect size estimates 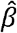 for cumulative flowering (in units of standard deviations, i.e., z-scores) ranged from −0.57 to 0.28 in the *S. dioica* habitat and from −0.52 to 0.34 in the *S. latifolia* habitat. Positive or negative allelic effects in one habitat (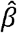 >> 0 or 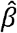 << 0) were associated with very low 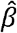 values (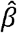 ≈ 0) in the other habitat, consistent with conditional neutrality (Fig. 3, cross pattern). A single locus showed opposing effects in the two habitats, 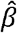 << 0 in the *S. dioica* habitat and 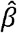 >> 0 in the *S. latifolia* habitat, suggestive of antagonistic pleiotropy (Fig. 3). Analyses for model-averaged effect sizes, 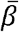, exhibited a very similar pattern (Supporting information Fig. S5), but with much smaller values due to multiplication with low posterior inclusion probabilities.

**Figure 3.**
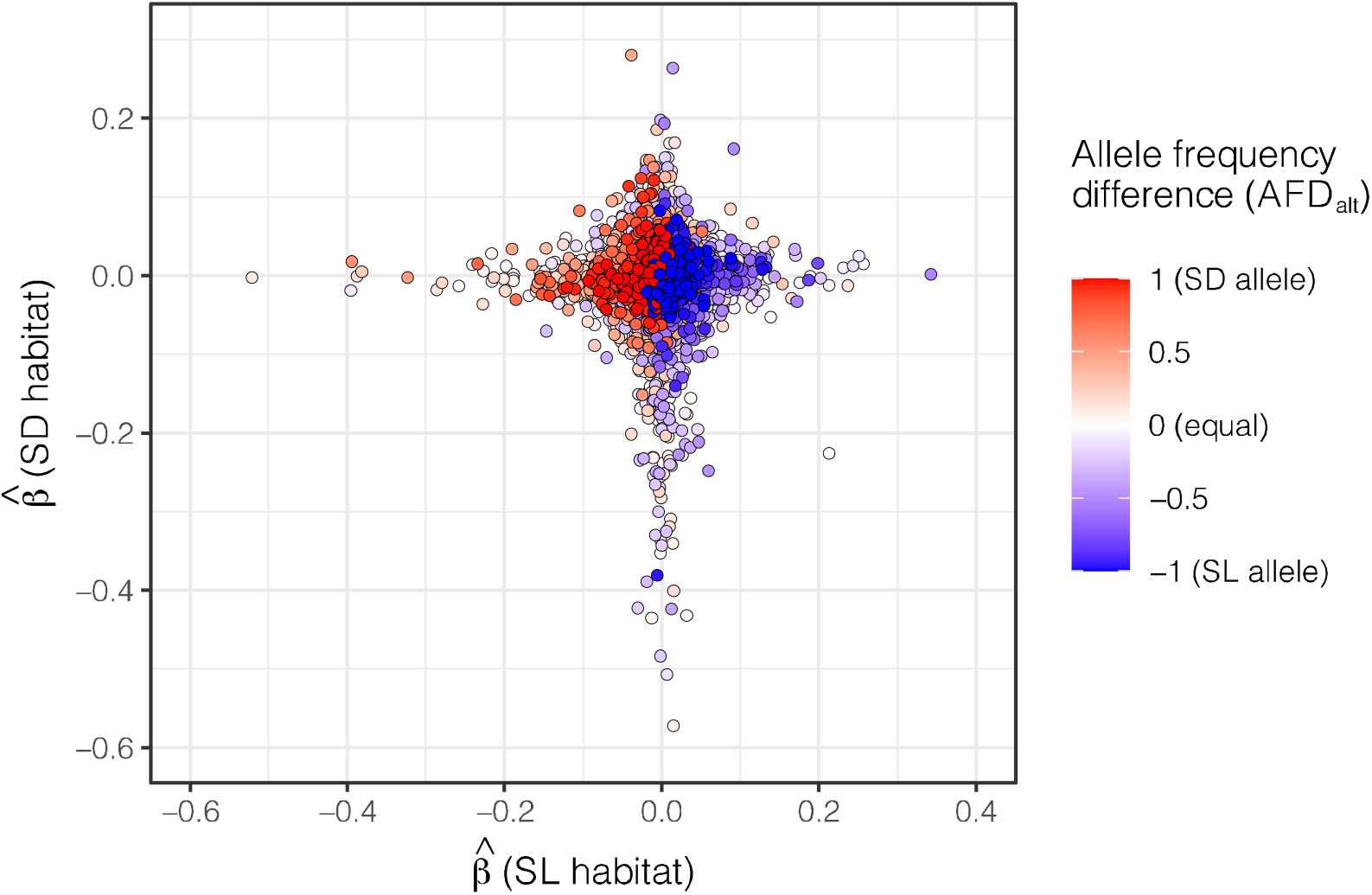
Locus-specific effect sizes of the alternative allele on cumulative flowering, a fitness component, measured in recombinant hybrids between two campion species, *Silene dioica* (SD) and *S. latifolia* (SL) that were transplanted into the habitat of each species (SD habitat, y-axis; SL habitat, x-axis). Given are raw effect estimates, 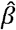, from Bayesian Sparse Linear Mixed Models (BSLMMs). Allele frequency differences for the alternative allele (AFD_alt_) between the two species (SD-SL) are indicated as a color gradient from blue (allele fixed in SL and absent in SD) to white (equal frequency in both species) to red (allele fixed in SD and absent in SL). Points are plotted in the order of increasing absolute AFD_alt_, such that highly differentiated loci are most visible. Loci with evidence for antagonistic pleiotropy would be have 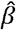-values of opposite signs in two habitats (top left, and bottom right), whereas loci consistent with conditional neutrality lie at zero on one axis, but deviate substantially from zero on the other.

A striking pattern in our data is that alleles with effects on cumulative flowering were significantly associated with higher allele frequencies in F_0_ individuals of the native species whereas alleles with negative effects on cumulative flowering were significantly more likely to be derived from the non-native species (Fig. 3, Supporting Information Figs. S5, S6 and S7). In the *S. dioica* habitat, loci with positive effects of the alternative allele on cumulative flowering had a positive median AFD_alt_ (alternative allele more common in *S. dioica*, red on Fig. 3), that were significantly different from the negative median AFD_alt_ (alternative allele more common in *S. latifolia*, blue on Fig. 3) at negative-effect alleles (Supporting Information Figs. S6 and S7). In the *S. latifolia* habitat, we observed the reverse pattern: loci with positive effects had negative median AFD_alt_ values that differed significantly from the positive median AFD_alt_ values for negative-effect loci (Supporting information Figs. S6 and S7). These patterns were observed for both 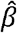 and 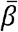, when using a subset of the approximately 0.5 % loci with strongest effect sizes (absolute values of 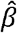 over 0.1 and absolute values of 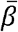 over 0.003) and when using all loci (i.e., negative-effect loci with 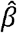 or 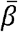 > 0 and positiveeffect loci with 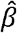 or 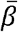 < 0, Supporting Information Figs. S6 and S7).

Alleles with positive effects on cumulative flowering in each habitat (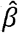 > 0.1) were rare in the foreign species but at appreciable frequencies in the native species; this effect was stronger in the *S. latifolia* habitat than in the *S. dioica* habitat (Fig. 4). Loci with negative effects (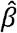 < −0.1), in contrast, were rare in the native species but at moderate or high frequencies in the foreign species (Fig. 4). These allele frequency patterns across species differ strongly from the overall allele joint allele frequency pattern for all loci (Fig. 2c).

**Figure 4.**
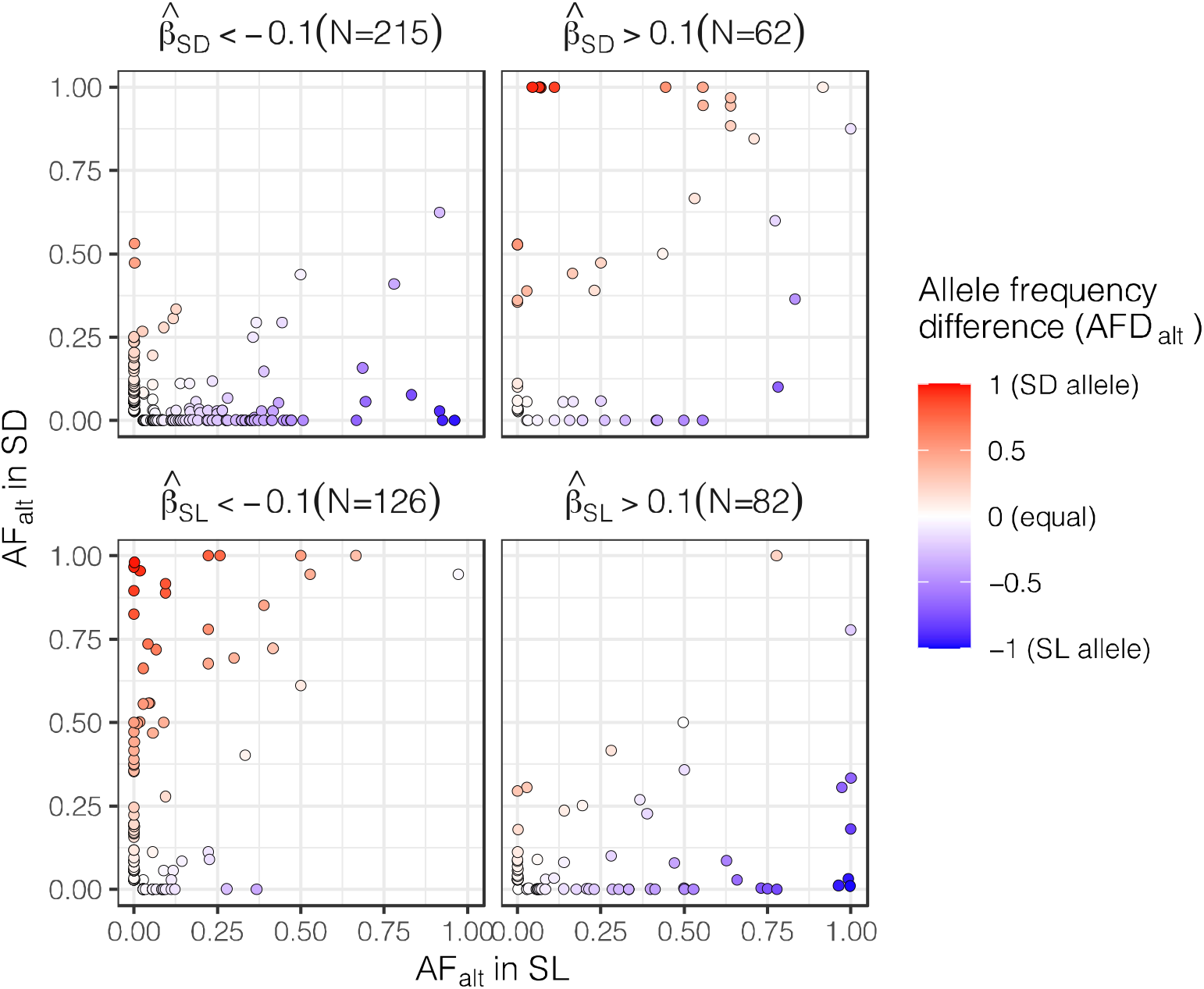
Frequencies of the alternative allele (AF_alt_) in two campion species, *Silene dioica* (SD, y-axis) and *S. latifolia* (SL, x-axis) for loci associated with cumulative flowering, a fitness component, in the habitat of each species. Shown are loci with absolute raw effect sizes, 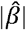, over 0.1 (in standard deviation units) in Bayesian Sparse Linear Mixed Models (BSLMMs) with the number of loci (N): 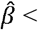 −0.1 in the SD habitat (negative selection), 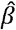 > 0.1 in the SD habitat (positive selection), 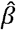 < −0.1 in the SL habitat (negative selection), and 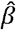 > 0.1 in the SL habitat (positive selection). Allele frequency differences for the alternative allele (AFD_alt_) between the two species (SD-SL) are indicated as a color gradient from blue (allele fixed in SL and absent in SD) to white (equal frequency in both species) to red (allele fixed in SD and absent in SL). Positive selection is associated with native alleles, whereas alleles under negative selection are more common in the non-native species (see also text and Supporting Information Figs. S6 and S7).

### Method validation

BSLMM estimates of narrow-sense heritability (*h^2^*) in simulated datasets responded to changes in simulated heritabilities, as expected (Fig. 5). BSLMMs on simulated data with a low *h^2^* of 0.05 recovered *h^2^* estimates in the correct range, while models on simulated data with a high *h^2^* of 0.2 yielded lower than expected *h^2^* estimates, especially when the number of sparse-effect loci was high (N = 50, Fig. 5). The number of sparse-effect loci was correctly estimated for simulated datasets with few loci (N = 10), but strongly underestimated when the simulated number of loci was high (Fig. 5). Correlations of simulated and estimated 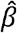 and 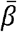-values were highest for datasets with high simulated heritabilities and declined for datasets with lower heritability and a higher number of functional loci (Fig. 5, Supporting Information Fig. S8). In simulations with *h^2^* = 0.02, 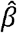 exhibited higher correlations with simulated effects sizes than 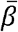 (Supporting Information Fig. S8).

**Figure 5.**
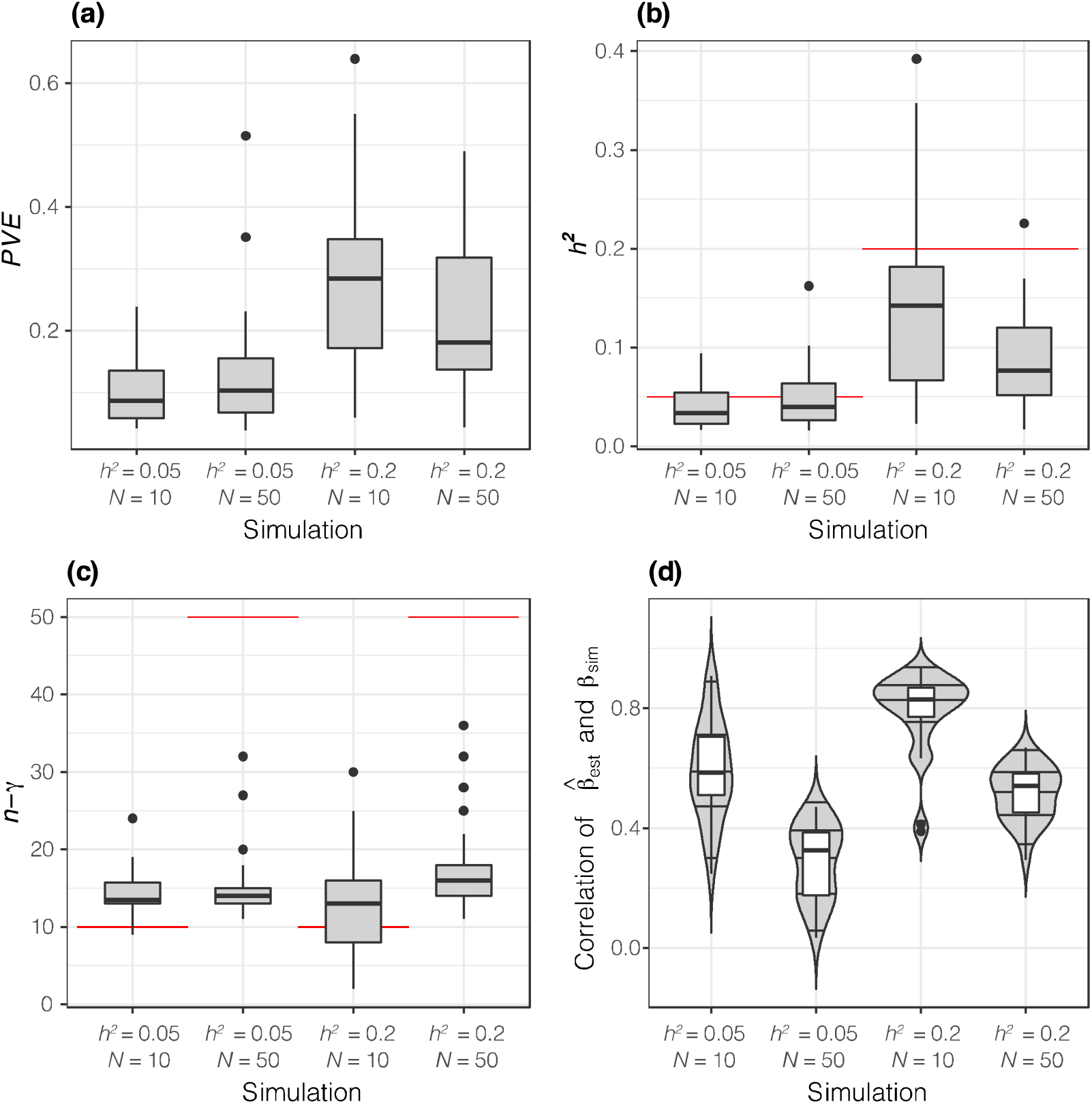
Performance analysis of Bayesian Sparse Linear Mixed Models (BSLMM) on simulated data based on a dataset from campions (*Silene*, 150 individuals, 42’090 loci). Simulated phenotypes had narrow-sense heritabilities (*h^2^*) of 0.05 or 0.20 with 10 or 50 functional loci (*N*); 30 replicate simulations per scenario were used and boxplots or violin plots summarize variation in medians of posterior distributions of hyperparameters among simulation replicates as well as of correlations of simulated and estimated per-locus effect sizes. **(a)** percent variation explained (*PVE*, interpreted as broad-sense heritability, *H^2^*), **(b)** narrow-sense heritability *h^2^* (as *PVE* * *PGE* [*PGE*, proportion of sparse effects in *PVE*]), **(c)** number of sparse-effect loci (*n-γ* and **(d)**, correlation of raw estimated effect sizes for individual loci 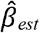 with simulated effect sizes *β_sim_*. On **(b)** and **(c)** simulated values are indicated with red lines.

These simulations thus show that our analyses can recover heritabilities and general patterns in effect size, however, estimates of the number of sparse effect loci (*n-γ*) must be treated with great caution. The consistency between estimated and simulated *h^2^* for datasets with low simulated *h^2^* shows that the low *h^2^* on our empirical data are reliable, especially as such low *h^2^* estimates were very rare in the high *h^2^* simulations. We expect that the apparent underestimation of *n-γ* also decreases posterior inclusion probabilities and thus 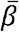.

## Discussion

Our results suggest that ecological differentiation between the campions *Silene dioica* and *S. latifolia* has a polygenic architecture, based on a reciprocal transplant experiment with recombinant second-generation hybrids between the two species. Many alleles had small effects on cumulative flowering, a fitness component, but only in one of the two habitats, conforming to a conditional neutrality pattern. Despite small effect sizes, the direction of allelic effects was consistent with selection for native alleles and against non-native alleles. This suggests that habitat adaptation is maintained by species-specific variants and that habitat dependent selection on hybrids reduces the inflow of non-native alleles. Below we discuss the implications of these findings for lineage divergence as well as the advantages and limitations of our approach.

### Low heritability of a fitness component in the field, based on many small-effect loci

Heritability of cumulative flowering, a fitness component, was low to moderate in F_2_ hybrids between *S. dioica* and *S. latifolia* in the habitats of both species. Using Bayesian Sparse Linear Mixed Models (BSLMM) we estimated narrow-sense heritability *h^2^* (*PVE* × *PGE*) to be 0.04 and 0.03 and broad-sense heritability *H^2^* (*PVE*) to be 0.12 and 0.10 for *S. dioica* and *S. latifolia* habitats, respectively. Our simulations indicate that such low *h^2^* values can be recovered fairly well with BSLMM analyses, as also reported by Gompert *et al*. (2017). Similar low to moderate heritabilities have previously been found for complex traits, for example for height in *Pinus albicaulis* (Lind *et al*., 2017) and in *Populus* (Bresadola *et al*., 2019) as well as in *Arabidopsis thaliana* accessions, where heritability for fitness components varied strongly between experimental sites (Exposito-Alonso *et al*., 2019). Our analyses suggest a polygenic genetic architecture of cumulative flowering in *Silene* with estimates of the number of sparse-effect loci estimated to 11 in the *S. dioica* habitat and to 16 in the *S. latifolia* habitat; however, our simulations indicate that the number of sparse effect loci in our dataset is difficult to estimate with BSLMM analyses, similar to results from Gompert et al. (2017). In many other systems, oligo- or polygenic control of fitness components was detected in natural settings, especially when the number of contributing loci was explicitly modelled (Lind *et al*., 2017; Bresadola *et al*., 2019; Exposito-Alonso *et al*., 2019). A polygenic genetic architecture of fitness components in our study system does appear likely given that QTL studies detected many loci distributed throughout the genome that were associated with ecologically relevant traits (Liu & Karrenberg, 2018; Baena□Díaz *et al*., 2019).

Limitations of studies on the genetic architecture of fitness in natural habitats include the number of individuals and sites that can be studied as well as the genetic resolution in the experimental material. Where markers are used, as in our study, most associations will be the result of linkage to causal variants rather than due to causal variants themselves. This reduces estimates of effect sizes and may lead to a situation where multiple loci linked to the same causal variant are included in alternative BSLMM iterations reducing the posterior inclusion probability (*PIP*) of each linked locus (Bresadola *et al*., 2019). We used a highly variable multi-cross F_2_ population with a limited number of recombination events, where markers likely were associated with larger genomic regions derived from each species. Recombination and genetic variation included in our material differs markedly from other studies using largely homozygous selfing accessions (*Arabidopsis*, Exposito-Alonso *et al*., 2019), within-species crosses or recombinant inbred lines derived from a small number of individuals (*Arabidopsis*, Fournier-Level *et al*., 2013; Leinonen *et al*., 2013; Ågren *et al*., 2017) or naturally recombinant wild-collected hybrids (*Populus*, Bresadola *et al*., 2019). These differences can make it difficult to compare results and model performance across studies. Despite its limitations, our setup has the advantage that the multi-cross F_2_ population used here approximates a natural contact site situation, where early-generation hybrids grow in the habitats of the two species when they come into contact (Karrenberg & Favre, 2008).

### Conditional neutrality is the prevailing pattern

Allelic effects conformed to a conditional neutrality pattern in our study: both positive and negative allelic effects on cumulative flowering in one habitat were associated with near-zero effects in the other habitat; only one locus deviated from this pattern. This finding further strengthens the view that conditional neutrality is more common in natural settings than antagonistic pleiotropy, as has been shown in monkeyflowers (Hall *et al*., 2010) and *Arabidopsis* (Fournier-Level *et al*., 2013; Ågren *et al*., 2017; Exposito-Alonso *et al*., 2019), switchgrass (Lowry *et al*., 2019) and *Lycaeides* butterflies (Gompert *et al*., 2015). However, strong evidence for conditional neutrality is generally difficult to provide, because this would require evidence for the absence of an effect in one of the habitats (Anderson *et al*., 2014; Mee & Yeaman, 2019). Interestingly, antagonistic pleiotropy appears to be detected more readily in controlled experimental evolution studies with microorganisms than in natural settings with higher plants or insects (Gompert & Messina, 2016; Bono *et al*., 2017; Wadgymar *et al*., 2017; Tusso *et al*., 2021). This is likely due to the control of selective agents in experiments as opposed to natural sites where both selective agents and traits under selection may vary (Wadgymar *et al*., 2017). In our system, high-elevation sites of *S. dioica* exhibit high winter mortality in *S. latifolia* and in hybrids in comparison to *S. dioica* (Favre *et al*., 2017) suggesting that frost tolerance or the regulation of carbohydrate storage could be under selection. At lowland *S. latifolia* sites, in contrast, *S. dioica* and hybrids suffer higher summer mortality than the native *S. latifolia*, most likely due to drought exposure (Favre *et al*., 2017). We therefore find it plausible that phenotypic and genetic targets of selection differ between habitats and this can contribute to the conditional neutrality pattern of effect sizes for fitness components as observed here.

### Fitness is increased by native alleles and decreased by non-native alleles, but not at the same loci

A striking result in our study is the association of allele frequencies in the two species with allelic effects on cumulative flowering, even for loci with very small effects. Alleles with positive effects were rare in the non-native species and common or of intermediate frequency in the native species, especially in the *S. latifolia* habitat. Alleles with positive effects on cumulative flowering in one habitat were neutral in the alternative habitat and could thus easily spread through both species if not lost by drift (Savolainen *et al*., 2013; Mee & Yeaman, 2019). Alleles with negative effects on cumulative flowering, on the other hand, had intermediate to high frequencies in the non-native species but were rare in the native species. Negative-effect alleles also included numerous variants that were rare in both species but these loci are most likely linked to causal loci with unknown allele frequencies rather than being causal loci themselves. Negative-effect alleles are most readily interpreted as deleterious load that is only exposed to selection in the alternative habitat and rare, conditionally deleterious variants are expected to arise frequently (Orr, 2005). Conditionally deleterious loci may experience reduced gene flow due to negative selection in the alternative habitat and thereby increase between-lineage differentiation (Mee & Yeaman, 2019). However, rare variants with conditionally deleterious effects likely remain rare even in the lineage they arose in and will thus not exhibit high differentiation between lineages. Variants with conditionally neutral effects, as reported here, likely play an important role for adaptation, but they are most certainly under-reported, not only because many of these variants are rare, but also because studies in alternative and relevant environments are needed to detect them.

### Implications for hybrid zones and speciation

Our results suggest that selection favors hybrids that are genetically similar to the native species. In early-generation hybrids with still large genomic regions of each species, as in our experiment, selection will favor genomic regions that contain positive-effect alleles that were more often derived from the native species. At the same time, selection will disfavor regions with negative-effect alleles that were often derived from the non-native species. Both types of conditionally neutral variants thus act as barriers to gene flow in each habitat. In natural hybrid zones between *S. dioica* and *S. latifolia*, intermediate individuals are rare and later-generation hybrids are heavily biased toward individuals that are genetically similar to the native species (Minder *et al*., 2007; Karrenberg & Favre, 2008). Our study suggests that this may be due not only to back-crossing with the native species but also to ecological selection in each habitat.

Regarding the evolution of adaptative differentiation, our results suggest adaptation through selection on many loci in each species. Alleles with small positive effects had intermediate to high frequencies in the native species and low frequencies in the non-native species. This may point to a redundant genetic control of adaptation as also suggested by a QTL study under controlled conditions in our study system (Liu & Karrenberg, 2018) and by studies in other systems, for example in *Drosophila* (Barghi *et al*., 2019). Polygenic architectures of adaptation with redundant effects are expected to be transient in time (Yeaman & Whitlock, 2011; Yeaman, 2015). Thus, even when ecological differentiation is a likely driver of divergence, as in our study system (Goulson & Jerrim, 1997; Karrenberg & Favre, 2008; Karrenberg *et al*., 2019), this may not necessarily manifest in strong, range-wide genetic differentiation at the underlying loci.

### Conclusion

Overall, our study adds to the understanding of how ecological differentiation can promote reproductive isolation and lineage divergence. We detected a polygenic architecture for a fitness component, cumulative flowering, in the campions *Silene dioica* and *S. latifolia* using a reciprocal transplant experiment. Allelic effects were mostly beneficial or deleterious in one habitat and neutral in the other and native-derived alleles were favored. Conditionally neutral effects may result from differences in selection targets in the two habitat types. Variants with conditionally neutral effects provide hidden variation upon which selection can act and facilitate current and future adaptation. The challenges ahead lie in understanding how within-lineage variation for fitness affecting alleles can be reconciled with the evolution of genome-wide differentiation between species.

## Acknowledgements

We thank to Rasmus Janson and Karin Steffen for providing help in the molecular lab. We are grateful for support from the Science for Life Laboratory (SciLifeLab) and the National Genomics Infrastructure, NGI, for massive parallel sequencing. Computations were performed on resources provided by SNIC through the Uppsala Multidisciplinary Center for Advanced Computational Science (UPPMAX) under SNIC projects 2017/7-406 and 2019/8-21. This work was funded by a project grants of the Swiss National Science Foundation (SNF, no. 3100A-118221), the Swedish Research Council (Vetenskapsrådet, no. 2012-03622) and the Carl Tryggers foundation to SK, as well as by a grant of the German Science Foundation (Deutsche Forschungsgemeinschaft, project no. FA1117/1-2) to AF.

## Author contribution

The study was designed by SK with input by AF and XL. AF conducted the transplant experiment with help from SK, XL performed molecular lab work, XL and SG did bioinformatic analyses, SG performed the association analysis and simulations with help from AB and SK. SK analyzed phenotypes and produced figures with help from SG. SK and SG wrote the paper with input from all other authors.

## Data availability

Double-digest RAD (ddRAD) sequencing data is available on NCBI’s Short Read Archive (SRA, SUB8324195 https://www.ncbi.nlm.nih.gov/sra/PRJNA669447). The variant call format (VCF) file as well as GEMMA input files will be submitted to dryad (project number to be added).

## Supporting Information

**Figure S1.** Principal components analysis including F_0_ and F_2_ generations.

**Figure S2.** Principal components analysis of F_2_ families.

**Figure S3.** Posterior distributions of hyperparameters, *S. dioica* habitat.

**Figure S4.** Posterior distributions of hyperparameters, *S. latifolia* habitat.

**Figure S5.** Model-averaged effect sizes 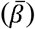 in the *S. dioica* habitat, plotted against 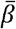 in the *S. latifolia* habitat.

**Figure S6.** Permutation test comparing between-species allele frequency differences among positive- and negative-effect alleles, based on raw effect size estimates 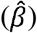.

**Figure S7.** Permutation test comparing between-species allele frequency differences among positive- and negative-effect alleles, based on model-averaged effect size estimates 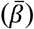.

**Figure S8.** Correlations among simulated effects and raw and model-averaged estimates of effect sizes (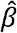 and 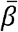).

**Table S1.** Description of source populations and transplant sites.

**Table S2.** Crossing scheme.

**Table S3.** Individuals used for crosses.

**Table S4.** Sparse-effect loci with posterior inclusion probabilities over 0.01.

