## Supporting Information Fig for "A polygenic architecture with conditionally neutral effects underlies ecological differentiation in *Silene*"

### Supplementary Figure S1

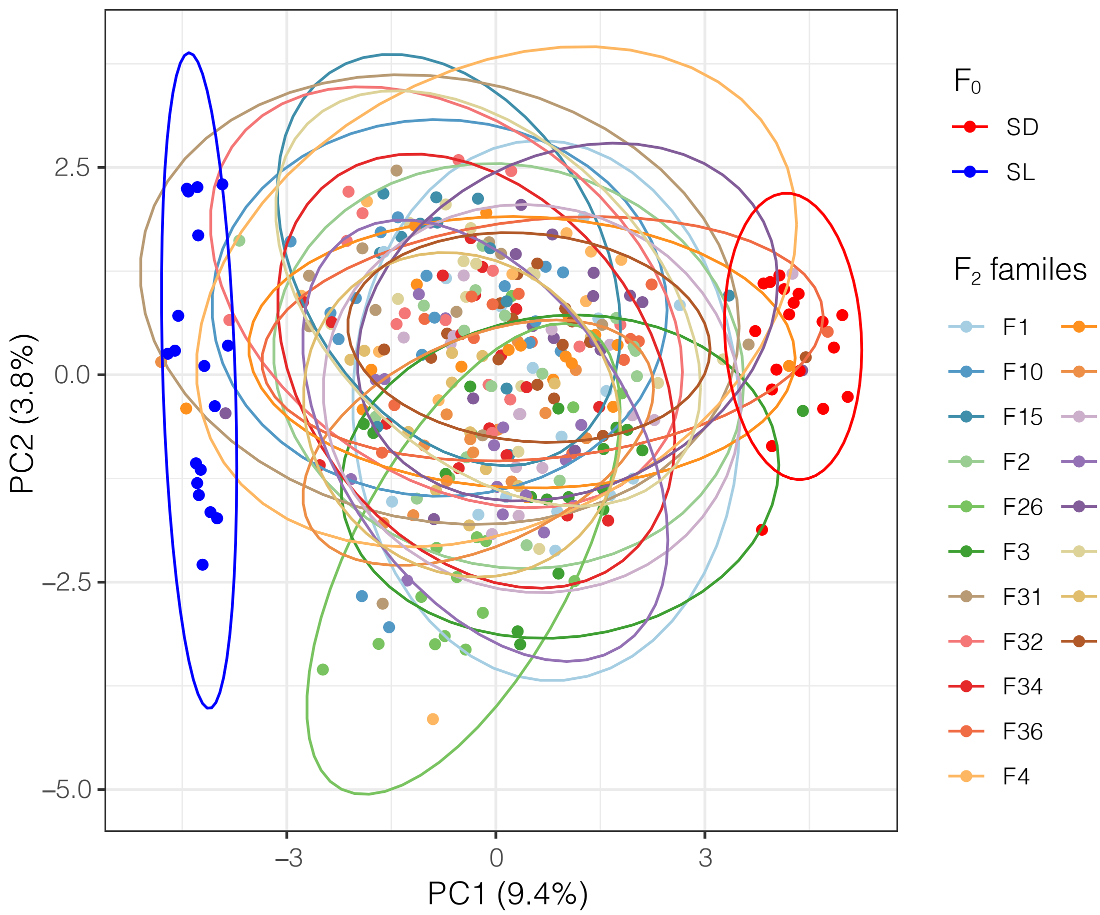


**Figure S1.** Principal component analysis of *Silene dioica* (SD, 18 individuals, $F_0$ generation) and *S. latifolia* (SL,18 individuals, $F_0$ generation) and of 19 families of second-generation hybrids ($F_2$, 290 individuals in total) between the two species, based on genotype probabilities at 220 SNP loci, for which all individuals had data. The first principal component explained 9.4% of the variation and the second principal component 3.8%. Analyses with more loci and imputed genotype probabilities for missing values did not improve the analysis.

**Supplementary Figure S2**

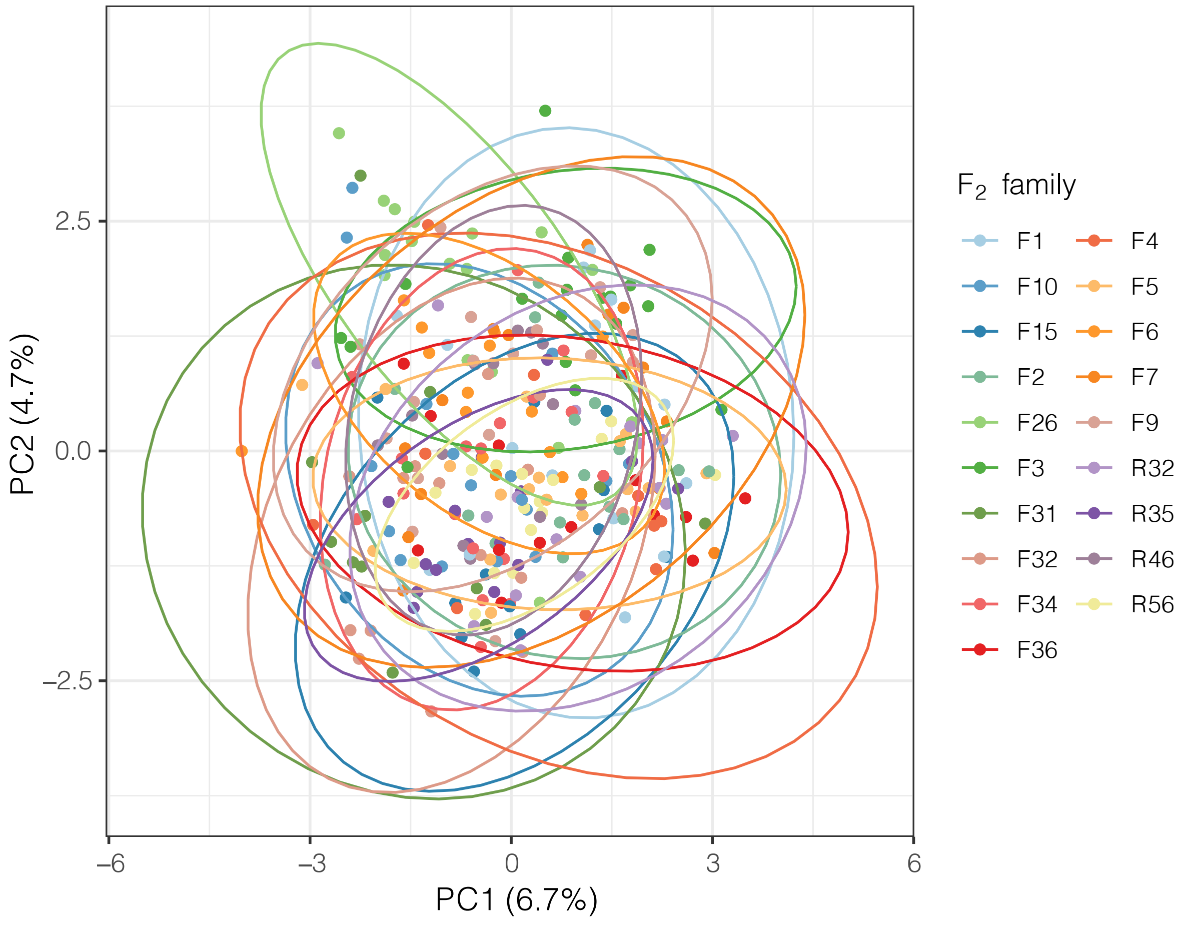


**Figure S2.** Principal component analysis of 19 families of second-generation hybrids ($F_2$, 290 individuals in total) between the campions *Silene dioica* and *S. latifolia*, based on genotype probabilities at 220 SNP loci, for which all individuals had data. The first principal component explained 6.7% of the variation and the second principal component 4.7%. Analyses with more loci and imputed genotype probabilities for missing values did not improve the analysis.

### Supplementary Figure S3

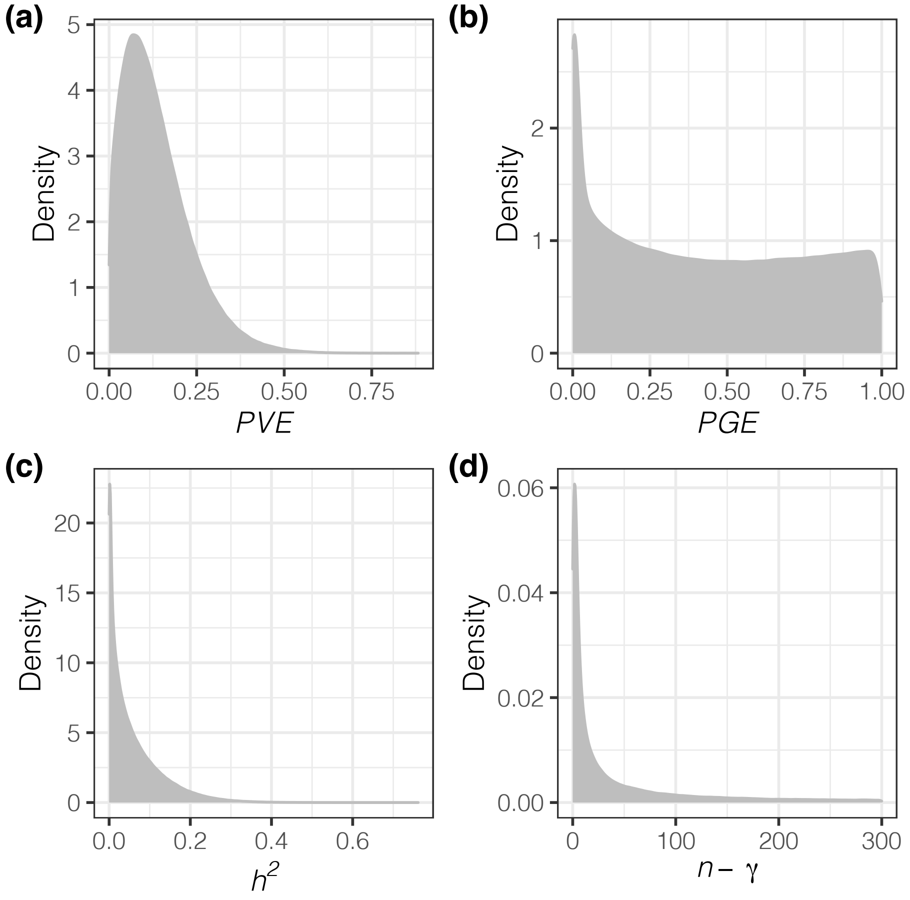


**Figure S3.** Posterior distributions of the hyperparameters, proportion of variation explained, *PVE* **(a)**, proportion of variation explained my loci with measurable effects, *PGE* **(b)**, narrow-sense heritability, $h^2$ **(c)** and the number of loci with measurable effects, $n$-$\gamma$ **(d)** in a genetic association analysis using a Bayesian Sparse Linear Mixed Model (BSLMM) of a fitness component, cumulative flowering, in second-generation hybrids between the campions *Silene dioica* and *S. latifolia* transplanted into the *S. dioica* habitat.

### Supplementary Figure S4

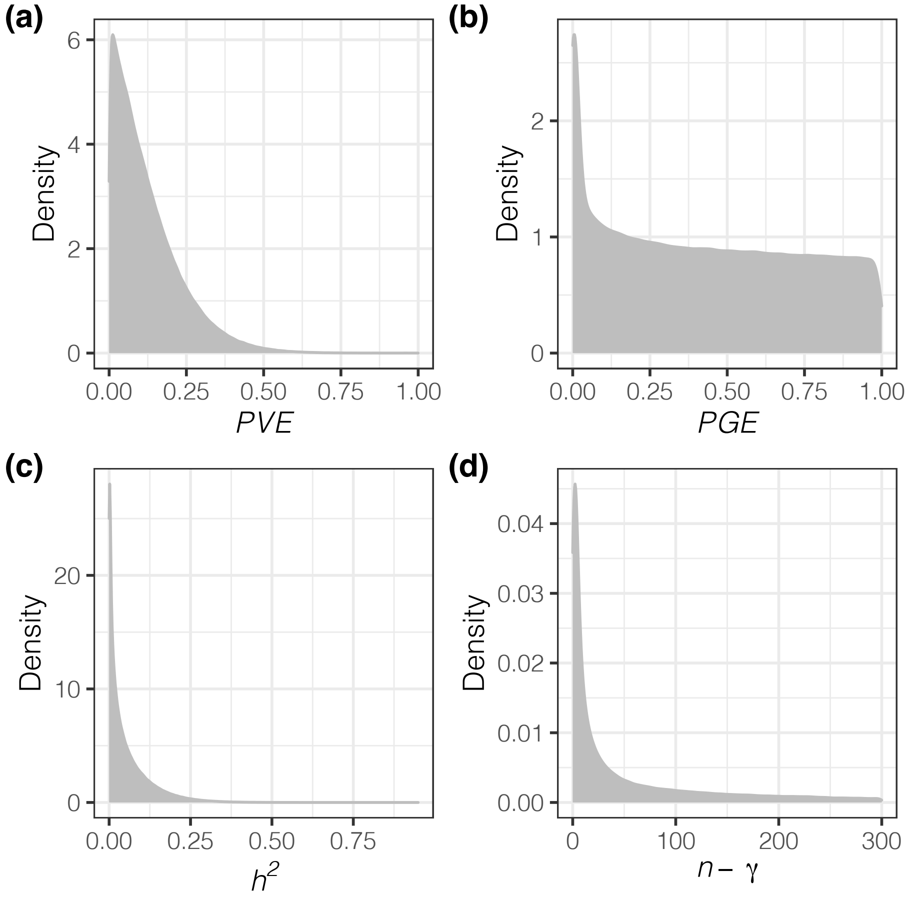


**Figure S4.** Posterior distributions of the hyperparameters, proportion of variation explained, *PVE* **(a)**, proportion of variation explained by loci with measurable effects, *PGE* **(b)**, narrow-sense heritability, $h^2$ **(c)** and the number of loci with measurable effects, $n$-$\gamma$ **(d)** in a genetic association analysis using a Bayesian Sparse Linear Mixed Model (BSLMM) of a fitness component, cumulative flowering, in second-generation hybrids between the campions *Silene dioica* and *S. latifolia* transplanted into the *S. latifolia* habitat.

### Supplementary Figure S5

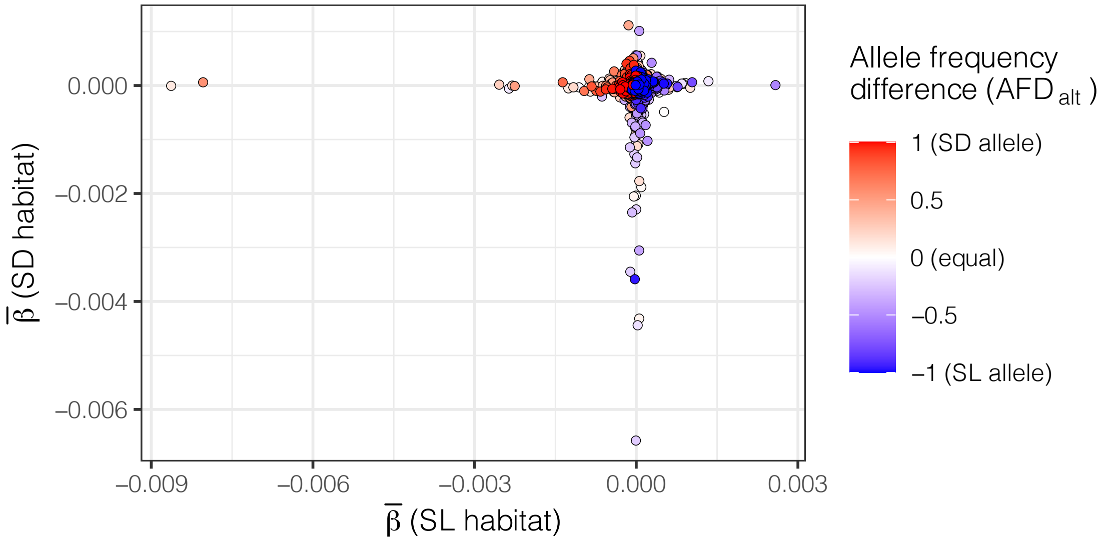


**Figure S5.** Locus-specific effect sizes of the alternative allele (model-averaged estimates, $\bar{\beta}$, from Bayesian Sparse Linear Mixed Models, BSLMMs) on cumulative flowering, a fitness component, measured in recombinant hybrids between two campion species, *Silene dioica* (SD) and *S. latifolia* (SL) that were transplanted into the habitat of each species (SD habitat, y-axis; SL habitat, x-axis). Allele frequency differences for the alternative allele ($AFD_{alt}$) between the two species (SD-SL) are indicated as a color gradient from blue (allele fixed in SL and absent in SD) to white (equal frequency in both species) to red (allele fixed in SD and absent in SL). Points are plotted in the order of increasing absolute $AFD_{alt}$, such that highly differentiated loci are most visible. Loci with evidence for antagonistic pleiotropy would be have $\bar{\beta}$-values of opposite signs in the two habitats (top left, and bottom right), whereas loci consistent with conditional neutrality lie at zero on one axis, but deviate substantially from zero on the other.

### Supplementary Figure S6

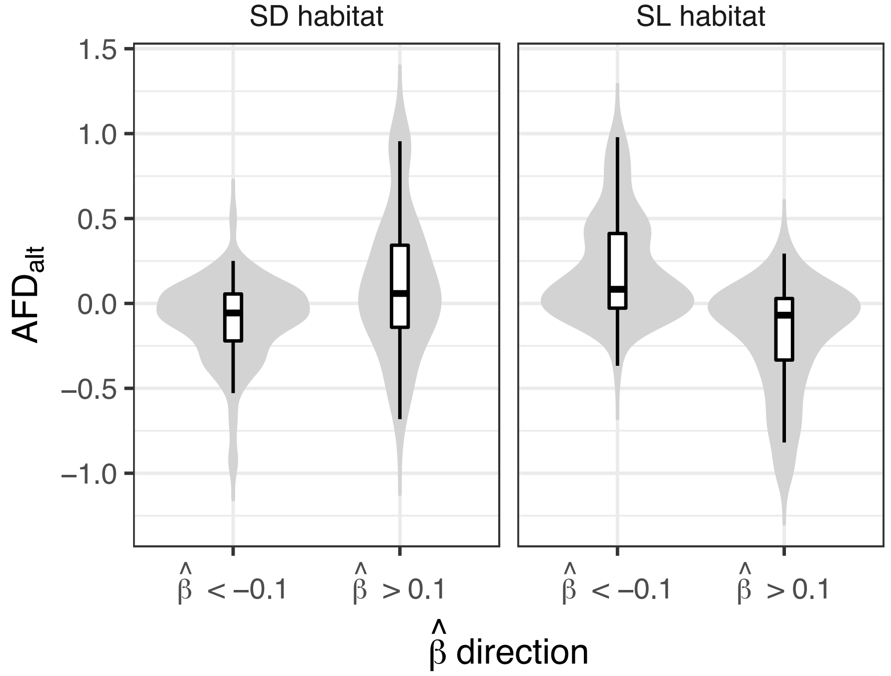


| | SD habitat | | SL habitat | |
|---|---|---|---|---|
| | $\hat{\beta}$ < -0.1 | $\hat{\beta}$ > 0.1 | $\hat{\beta}$ < -0.1 | $\hat{\beta}$ > 0.1 |
| N | 216 | 62 | 126 | 82 |
| Median | -0.056 | 0.058 | 0.083 | -0.069 |
| **Permutation test** | Z = -4.529 | | Z = 7.199 | |
| | P = 5.93 * $10^{-6}$ | | P = 6.08 * $10^{-13}$ | |

**Figure S6.** Permutation test comparing allele frequency differences ($AFD_{alt}$, polarized to the alternative allele) between the campions *Silene dioica* (SD) and *S. latifolia* (SL) at loci with negative ($\hat{\beta}$ < -0.1) and positive effects ($\hat{\beta}$ > 0.1) on a fitness component, cumulative flowering, measured in second-generation hybrids between the two species transplanted to the habitat of each species (SD habitat and SL habitat). High $AFD_{alt}$ values indicate that the alternative allele is more common in SD and low values indicate that it is more common in SL. Effects sizes are based on raw effect estimates ($\hat{\beta}$) from Bayesian Sparse Linear Mixed Models (BSLMMs). Similar results were obtained when comparing loci with $\hat{\beta}$ < 0 and $\hat{\beta}$ > 0 (SD: Z = -6.440, *P* = 1.19 *$10^{-10}$; SL: Z = 29.180, *P* = < 2.2* $10^{-16}$).

### Supplementary Figure S7

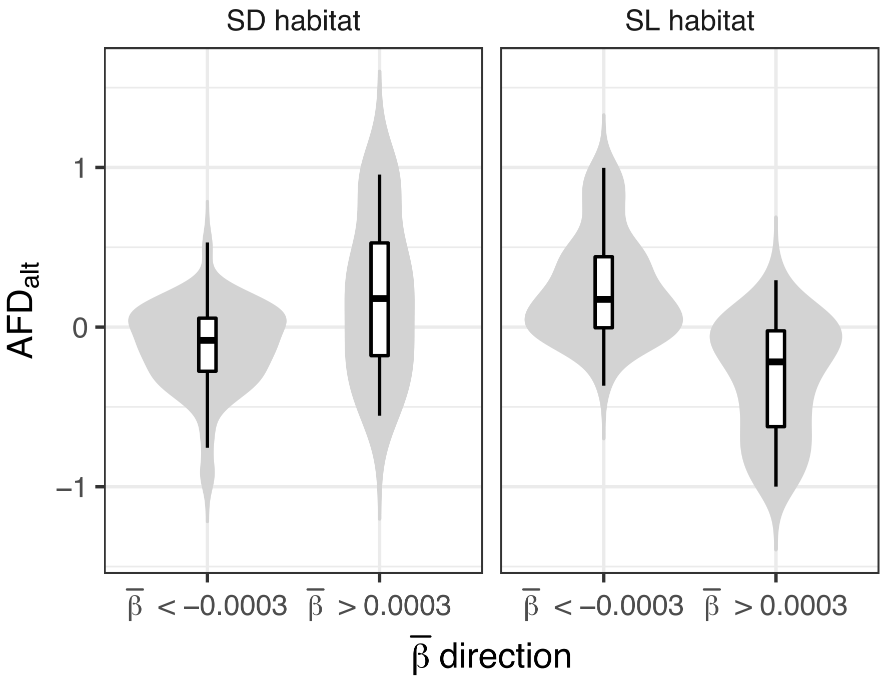


| | SD habitat | | SL habitat | |
|---|---|---|---|---|
| | $\bar{\beta}$ < -0.0003 | $\bar{\beta}$ > 0.0003 | $\bar{\beta}$ < -0.0003 | $\bar{\beta}$ > 0.0003 |
| N | 140 | 38 | 140 | 38 |
| Median | -0.083 | 0.179 | 0.173 | -0.218 |
| **Permutation** | Z = -4.511 | | Z = 9.090 | |
| **test** | $P = 6.46 * 10^{-6}$ | | $P = <2.20 * 10^{-16}$ | |

**Figure S7.** Permutation test comparing allele frequency differences ($AFD_{alt}$, polarized to the alternative allele) between the campions *Silene dioica* (SD) and *S. latifolia* (SL) at loci with negative ($\bar{\beta}$< -0.0003) and positive effects ($\bar{\beta}$ > 0.0003) on a fitness component, cumulative flowering, measured in second-generation hybrids between the two species transplanted to the habitat of each species (SD habitat and SL habitat). High $AFD_{alt}$ values indicate that the alternative allele is more common in SD and low values indicate that it is more common in SL. Effects sizes are based on model-averaged effect estimates ($\bar{\beta}$) from Bayesian Sparse Linear Mixed Models (BSLMMs). Similar results were obtained when comparing loci with $\bar{\beta}$ < 0 and $\bar{\beta}$ > 0 (SD: Z = -6.443, $P = 1.17 * 10^{-10}$; SL: Z = 28.212, $P = < 2.20 * 10^{-16}$).

### Supplementary Figure S8

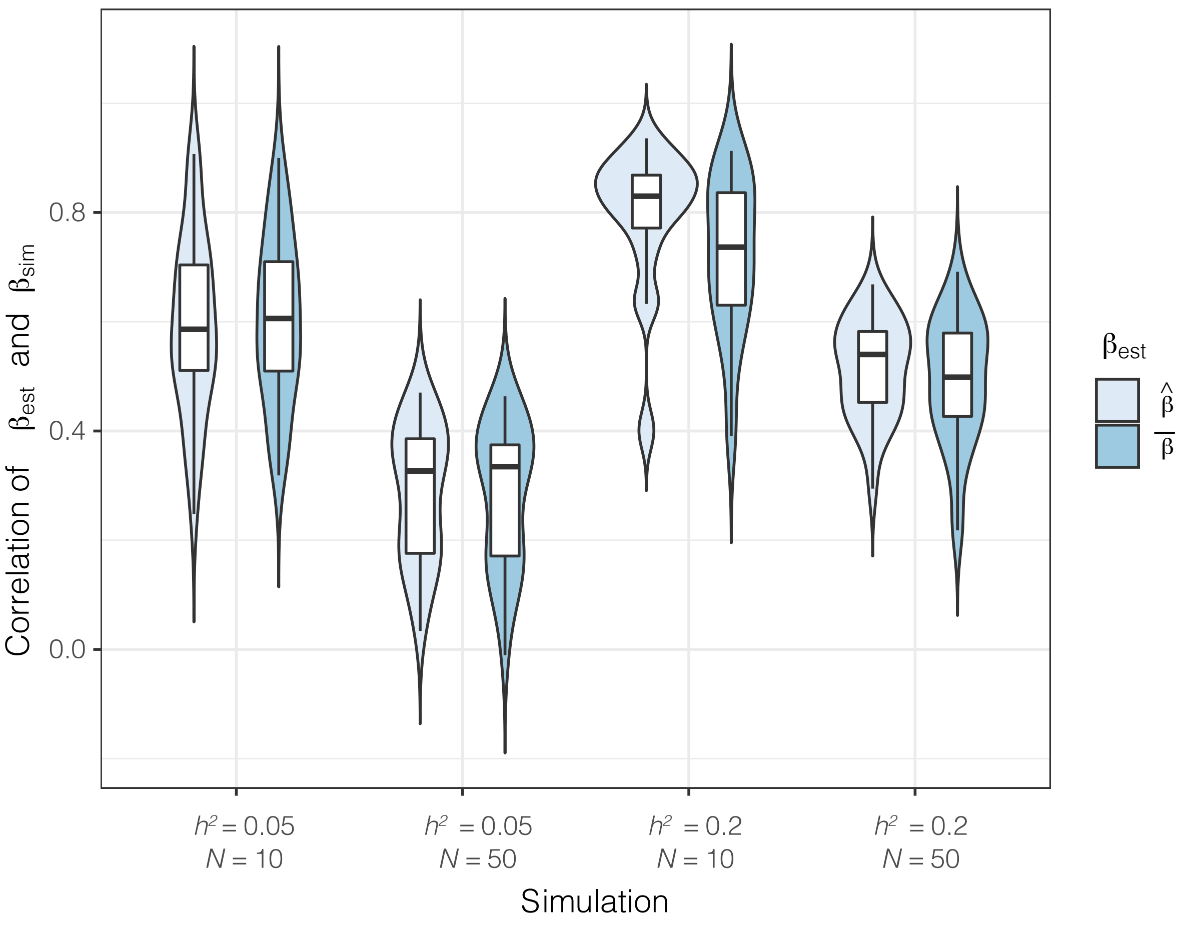


**Figure S8.** Correlations among simulated effects and raw and model-averaged estimates of effect sizes ($\hat{\beta}$ and $\bar{\beta}$, respectively) for a genetic association analysis with Bayesian Sparse Linear Mixed Models (BSLMMs) on a dataset from campions (*Silene*, 150 individuals, 42'090 loci). Raw effect sizes, $\hat{\beta}$, are estimated assuming $\beta > 0$, while model-averaged effect sizes, $\bar{\beta}$, take posterior inclusion probabilities into account. Simulated phenotypes had narrow-sense heritabilities ($h^2$) of 0.05 or 0.20 with 10 or 50 functional loci (*N*); 30 replicate simulations per scenario were used and boxplots or violin plots summarize variation among simulation replicates.
